# A potent synthetic nanobody with broad-spectrum activity neutralizes SARS-Cov-2 virus and Omicron variant through a unique binding mode

**DOI:** 10.1101/2022.04.11.487660

**Authors:** Dongping Zhao, Liqin Liu, Xinlin Liu, Jinlei Zhang, Yuqing Yin, Linli Luan, Dingwen Jiang, Xiong Yang, Lei Li, Hualong Xiong, Dongming Xing, Qingbing Zheng, Ningshao Xia, Yuyong Tao, Shaowei Li, Haiming Huang

**Author notes:** To whom correspondence should be addressed. E-mail: Haiming Huang, Shaowei Li(.), Yuyong Tao. These authors contributed equally to this work (Dongping Zhao, Liqin Liu, Xinlin Liu, Jinlei Zhang).

## Abstract

The major challenge to control COVID pandemic is the rapid mutation rate of the SARS-Cov-2 virus, leading to the escape of the protection of vaccines and most of the neutralizing antibodies to date. Thus, it is essential to develop neutralizing antibodies with broad-spectrum activity targeting multiple SARS-Cov-2 variants. Here, we reported a synthetic nanobody (named C5G2) obtianed by phage display and subsequent antibody engineering. C5G2 has a single digit nanomolar binding affinity to RBD domain and inhibits its binding to ACE2 with an IC_50_ of 3.7 nM. Pseudovirus assay indicated that the monovalent C5G2 could protect the cells from the infection of SARS-Cov-2 wild type virus and most of the virus of concern, i.e. Alpha, Beta, Gamma and Omicron variants. Strikingly, C5G2 has the highest potency against Omicron among all the variants with the IC_50_ of 4.9ng/mL. The Cryo-EM structure of C5G2 in complex with the Spike trimer showed that C5G2 bind to RBD mainly through its CDR3 at a conserved region that not overlapping with the ACE2 binding surface. Additionally, C5G2 bind simultaneously to the neighboring NTD domain of spike trimer through the same CDR3 loop, which may further increase its potency against the virus infection. Third, the steric hindrance caused by FR2 of C5G2 could inhibit the binding of ACE2 to RBD as well. Thus, this triple-function nanobody may be served as an effective drug for the prophylaxis and therapy against Omicron as well as future variants.

## Introduction

COVID-19 (coronavirus disease 2019) has became a pandemic worldwide since the end of 2019. As of Feb 25^th^ 2022, it has caused more than 420 million infection cases and more than 6 million deaths(https://covid19.who.int/). It is SARS-CoV-2, which belongs to the beta coronavirus family, that cause the disease[1, 2]. SARS-CoV-2 virus enter human cells via its spike protein, which is capable of binding to the ACE2(angiotensin-converting enzyme 2) receptors on the surface of the host cells[3, 4]. The spike protein is composed of S1 subunit at its N-termini and S2 subunit at its C-termini[5]. The S1 contains a N-terminal domain (NTD) and a receptor binding domain (RBD) which engages the binding to the ACE2 receptors of the host cells[5–7]. Upon the binding, S2 subunit will mediate the membrane fusion between the viral and host cells to finish the viral entry[8, 9]. Based on this biological process, spike protein has been the target for neutralizing antibody therapy and vaccine protection[10–12]. However, the rapid mutation rate of SARS-CoV-2 challenges the development of neutralizing antibody and vaccine.

There are two groups of neutralizing antibodies targeting spike protein of SARS-CoV-2, the majority of which are those binding to RBD, thus blocking virus interaction with ACE2[10, 11, 13]. In addition, antibodies targeting NTD could also have neutralization efficacy[14, 15]. The mechanism of anti-RBD neutralizing antibodies could also be classified into two groups[16]. The first group of antibodies bind to RBD and blocked its interaction with ACE2 directly, thus protecting the host cells from virus infection[10, 17, 18]. On the contrary, the other group of antibodies interact with RBD at epitopes distinct from the binding interface between RBD and ACE2, resulting virus entry prevention as well, due to partially, if not all, the conformation changes of RBD[19, 20]. This type of neutralizing antibodies is more likely to act as the broad-spectrum antibodies as they bind to the conserved residues at RBD/NTD across the SARS-CoV-2 variants.

The neutralizing antibodies were mainly derived from B cells of COVID-19 patients and immunized animals[21, 22] or phage displayed antibody libraries, either naturally origin or synthetic ones[10, 11, 17]. The synthetic antibody library technology harnesses the frameworks of natural antibodies, based on which only the CDRs(Complementarity-determining regions) were partially or completely randomized to form large content(diversity usually greater than 10^9^ pfu) libraries[23–25]. Though independent of immune system, the notion of synthetic antibody as therapeutic has been proved feasible and practical as exemplified by MorphoSys antibody drug guselkumab. The guselkumab, targeting IL23, was developed from MorphoSys synthetic library HuCAL and approved to treat plaque psoriasis by FDA in 2017[26, 27]. HuCAL library was constructed using human germlines as scaffolds[28]. Since then, many different antibody scaffolds have been employed to generate synthetic antibody libraries, including nanobodies. Nanobodies or single domain antibodies are the variable regions of the heavy-chain only antibodies that from Camelid immune system[29, 30]. One of the benefits of using nanobodies to construct synthetic antibody libraries is that a nanobody has only three CDRs(*versus* six CDRs in a conventional antibody) , which makes the mutation process easier than those of using conventional IgG(Fab or scFv fragment) as scaffolds[31]. There are many nanobodies, either natural[11, 32] or synthetic ones[10, 33], have been reported that have neutralizing activity against SARS-CoV-2 virus. Unfortunately, there are few nanobodies reported to date that have broad-spectrum neutralizing activities against multiple SARS-CoV-2 variants, especially the newly emerged Omicron.

We firstly described in this study the construction of a synthetic nanobody library based on a camel nanobody by a novel randomization strategy. We selected this library against SARS-CoV-2 spike protein, S1 subunit and RBD domain, respectively by phage display and obtained seven nanobodies in total. One of the RBD binding nanoboides, named C5, is capable of competing with ACE2 to bind to RBD. Protein engineering technology was applied to improve the binding affinity of C5 to RBD and its solubility and yield. The engineered variant, i.e. C5G2, has been proved its potency to protect living cells from infection by SARS-CoV-2 pseudovirus. In addition, C5G2 could inhibit multiple variants, including the Omicron, infecting living cells with high potency in pseudovirus assay. We further solved the cryo-EM structure of C5G2 in complex with the spike protein trimer, uncovering the structural basis of the broad-spectrum neutralizing activity.

## Results

### The design, construction and quality control of a synthetic nanobody library

In order to obtain antibodies for diverse antigens by phage display, we firstly sought to construct a synthetic nanobody library. To this end, a natural nanobody (PDB no. 1ZVH) [34], which recognizes lysozyme, was selected as the template for the library construction (Figure 1A). In our library design, residues from 27 to 33 of complementarity-determining region 1 (CDR1), 51 to 58 of CDR2 and 99 to 112 of CDR3 were randomized simultaneously (Figure 1A). The length of each CDR in the library were kept the same as the template. At each position of the CDRs, an amino acid mix were incorporated by Kunkel method[35, 36]. The amino acid mix(X) is composed of 19 natural amino acids, excluding cysteine (Figure 1A) to prevent the forming of unexpected disulfide bond though extra interloop disulfide bond is very common in the natural occurring nanobodies, especially camel derived ones [29] . The detail of the construction of the library was described in the Material&Methods section. The resultant diversity of the library was ∼3×10^9^ colony-forming units (cfu). The DNA deep sequencing result indicated that the mutated molecules have expected amino acid percentage as designed at each CDR (Figure 1B). This library was then named <u>A</u>UAM <u>sy</u>nthetic <u>na</u>nobody <u>l</u>ibrary (ASyNAL). We then panned the library against seven diverse antigens by phage display. Other than lysozyme, those antigens were cytoplasmic proteins i.e. SH2 domain of Tyrosine-protein kinase Fyn (Fyn_SH2) and Glutathione S-transferase (GST), extracellular domain of cell membrane proteins i.e. Human Epidermal Growth Factor Receptor 2 (Her2), Siglec15 and virus proteins i.e. human SARS-CoV-2 spike protein and Porcine Circovirus (PCV) 2d protein. As shown in Table 1, we were able to obtain at least one nanobody for each of those seven antigens verified by phage ELISA, indicating the versatile applications of ASyNAL in generation of nanobodies for diverse antigens.

**Figure 1.**
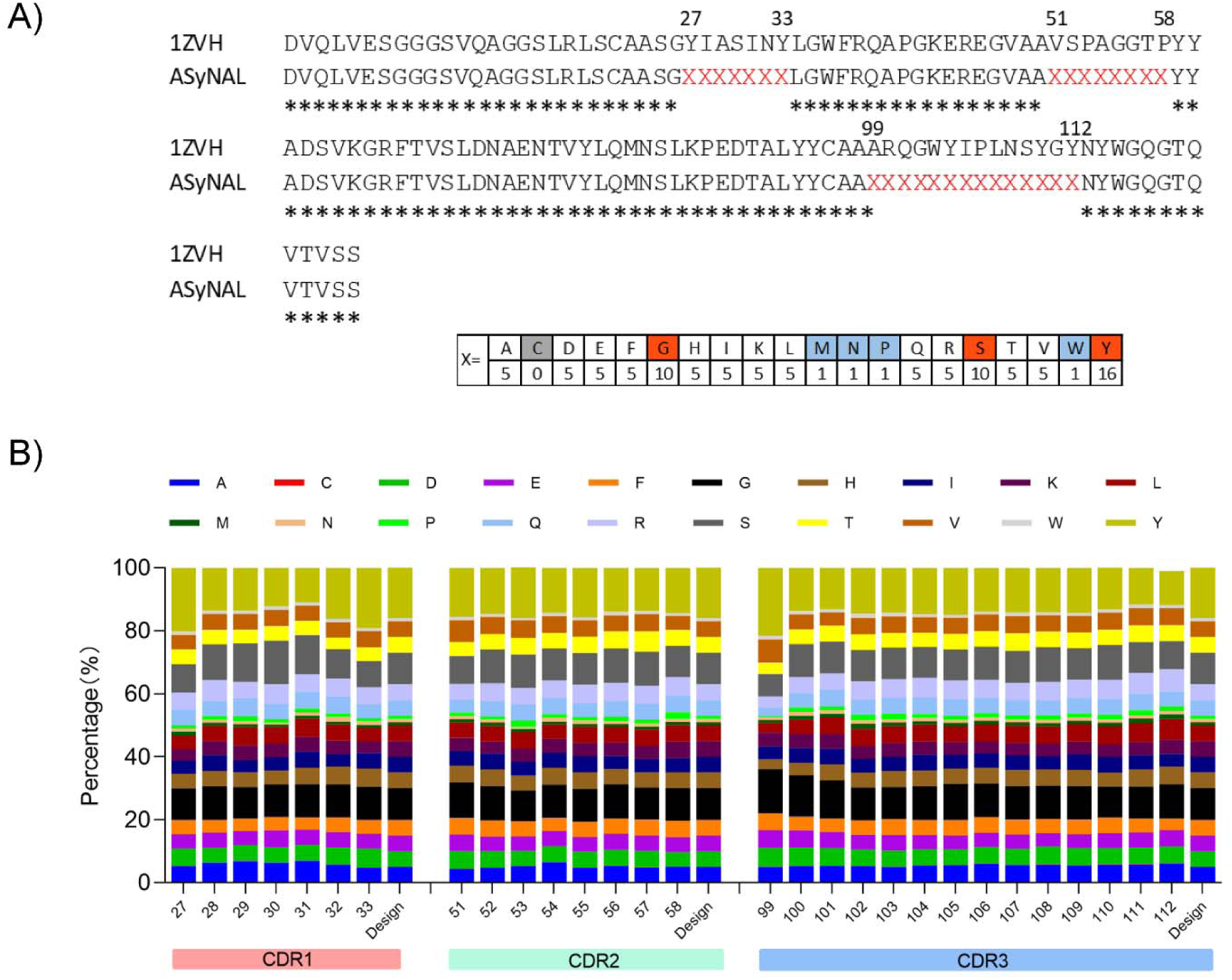
Construction and quality control of a synthetic nanobody library. A). Sequence alignment of a camel nanobody(1ZVH) with ASyNAL scaffold, showing the randomization regions in CDR1(27-33), CDR2(51-58)&CDR3(99-112) by X, respectively. The X is a mix of amino acids with the ratio showed below. B). NGS data(n=20112) showing the amino acid occurrence rate of the ASyNAL at each position *vs* designed rate. 20112 nanobody sequences were aligned, the amino acid residues at CDR regions were retrieved and the rate of each amino acid occurrence (%) at each position were calculated.

**Table 1.** Antigens and their nanobodies selected from ASyNAL library

| Antigen | Unique Nb | Category |
| --- | --- | --- |
| Lysozyme | 10 | Secreted enzyme |
| Fyn SH2 | 1 | Cytoplasmic protein |
| GST | 6 | Cytoplasmic protein |
| Her2 | 9 | Extracellular domain of membrane |
| Siglec 15 | 11 | Extracellular domain of membrane |
| SARS-CoV-2 spike | 7 | Virus protein |
| Swine Pcv-2d | 8 | Virus protein |

### Characterization of potential neutralizing nanobody against SARS-CoV-2 spike protein

In an attempt to obtain neutralizing nanobody against SARS-CoV-2, we panned the ASyNAL library against three antigens including the spike protein, S1 domain and RBD, respectively. As a result, we successfully obtained six spike-specific nanobodies (named C5, D3, D8, D9, F12 and G6), one S1-specific nanobody (A11) and one RBD-specific nanobody (B7) (Figure 2A-C, Table S1, Figure S1). DNA sequencing showed that D3 clone (from spike panning) and A11 (from S1 panning) are sequentially identical. Thus, we totally got seven unique nanobodies for spike protein. To map the binding epitopes of those nanobodies, we then applied cross-reactive phage ELISA assay to check the bindings of those seven nanobodies to spike protein and its fragments, respectively. All seven nanobodies bound the spike protein as expected (Figure 2A). Furthermore, they all bound to S1 as well (Figure 2B), but only nanobodies B7 and C5 could bind to the RBD (Figure 2C), implying the other five clones are likely NTD binders (Figure S1A). In a competitive ELISA, NTD binding nanobodies D8, D9, F12 and G6, but not RBD binder B7, inhibit D3 phage binding to spike protein (Figure S1B), showing they may share the same (or overlapping) epitope as D3 clone in the NTD domain (Figure S1A&Table S1).

**Figure 2.**
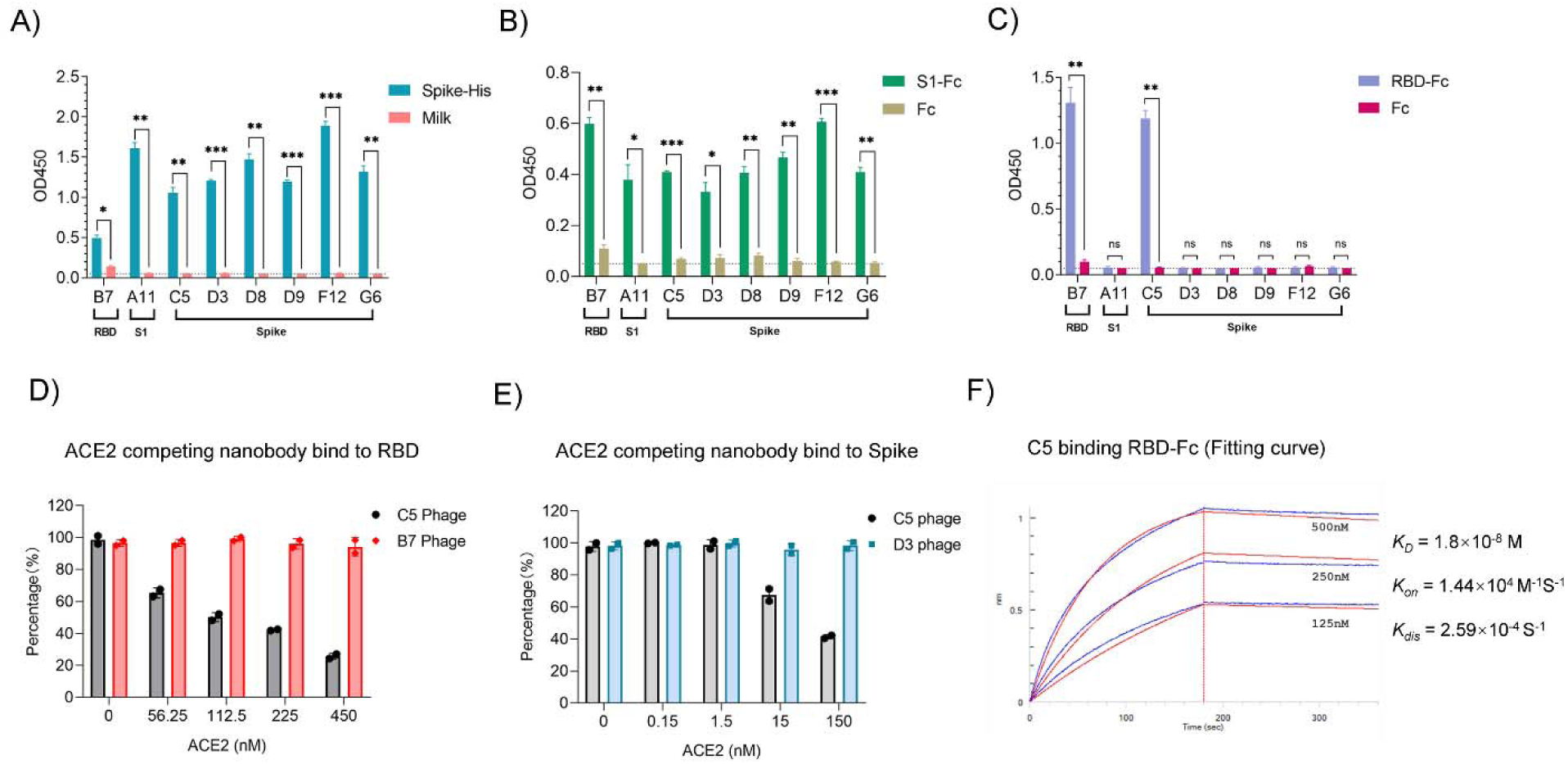
Identification of C5 clone as a spike protein/RBD binder and compete with both Spike and RBD binding to ACE2. Phage panning of the ASyNAL library against antigens RBD, S1 and spike protein respectively generated eight nanobody clones. Clone B7 is for RBD, A11 is for S1 and C5, D3, D8, D9, F12, G6 are for spike protein. The cross reactivities of the eight clones to Spike(A), S1(B) and RBD(C) were determined by phage ELISA. Two replicates were performed, and the p-value was performed with Student’s t-test (ns, no significant; *p < 0.05; **p<0.01; ***p<0.001). Competitive ELISA showed that C5 phage and ACE2 competed each other to bind RBD(D) and the full length spike protein(E). As controls, B7 phage and D3 phage had no competition. F) Binding curve of C5 protein to RBD in BLI assay. The affinity is 18 nM.

The binding affinities of those nanobodies to their screening antigens (spike, S1 or RBD) were measured by Bio-Layer Interferometry (BLI). As shown in Table 2, the binding affinities (KD) are ranging from 10^-10^ M to 10^-7^ M, which are similar to those selected from synthetic nanobody libraries[10] as well as natural nanobody libraries[11]. Furthermore, we investigated the size exclusion chromatography (SEC) profiles of the nanobodies and the result indicated that, other than C5 which has aggregation (data not shown), five nanobodies (B7, D8, D9, F12, G6) showed one single peak and nanobody A11 and D3 showed minimal (∼5%) aggregation (Figure S2). As the spike protein of SARS-CoV-2 binds to human Angiotensin-converting enzyme 2 (ACE2) through its RBD domain to initiate the entry of the virus into human cells, we wonder if the RBD-specific nanobodies B7 and C5 identified in this study could interfere with the binding of ACE2 to the RBD. Unfortunately, the yield of C5 protein was very poor when expressed in the bacterial host BL21. So we were unable to use C5 protein as competitor to interfere with the binding of spike protein to the ACE2 by competitive ELISA. Oppositely, we tried to use ACE2 to interfere with the binding of C5/B7 nanobodies displayed on phage to S protein and RBD (i.e. phage competitive ELISA). As shown in Figure 2D, the binding of C5, but not B7, to RBD could be inhibited when using ACE2 protein as the competitor. Furthermore, the binding of C5 to spike protein could also be blocked by ACE2 protein (Figure 2E). Based on the data, we deduced that C5 could be used as a blocker against spike binding to ACE2 to neutralize the virus.

**Table 2.** Anti-Spike nanobody affinity and neutralization potency

|  |  | B7 | A11/D3 | D8 | D9 | F12 | G6 | C5 | C5G2 | mNb6 |
| --- | --- | --- | --- | --- | --- | --- | --- | --- | --- | --- |
| Antigen |  | RBD | S1/Spike | Spike | Spike | Spike | Spike | Spike | Spike | Spike |
| Spike binding | KD(M) |  |  | 3.05E-08 | 2.61E-07 | 1.46E-07 | 2.48E-07 |  |  |  |
|  | Kon(1/Ms) |  |  | 7.71E+04 | 1.87E+05 | 2.51E+05 | 1.40E+05 |  |  |  |
|  | Kdis(1/s) |  |  | 2.53E-03 | 4.87E-02 | 3.66E-02 | 3.46E-02 |  |  |  |
| S1 binding | KD(M) |  | 5.58E-10 |  |  |  |  |  |  |  |
|  | Kon(1/Ms) |  | 8.83E+05 |  |  |  |  |  |  |  |
|  | Kdis(1/s) |  | 4.93E-04 |  |  |  |  |  |  |  |
| RBD binding | KD(M) | 2.66E-07 |  |  |  |  |  | 1.80E-08 | 1.62E-09 | 3.48E-09 |
|  | Kon(1/Ms) | 2.59E+05 |  |  |  |  |  | 1.44E+04 | 1.76E+04 | 1.63E+04 |
|  | Kdis(1/s) | 6.87E-02 |  |  |  |  |  | 2.59E-04 | 2.84E-05 | 5.66E-05 |
| Pseudo-virus IC<sub>50</sub>(M) | WT |  | N.E. |  |  |  |  | N.E. | 1.59E-09 | 1.37E-08 |
|  | Alpha |  | N.E. |  |  |  |  | N.E. | 9.5E-10 | 2.69E-08 |
|  | Beta |  | N.E. |  |  |  |  | 8.16E-08 | 5.6E-10 | N.E. |
|  | Gamma |  | N.E. |  |  |  |  | N.E. | 1.06E-09 | N.E. |
|  | Delta |  | N.E. |  |  |  |  | N.E. | N.E. | 2.05E-08 |
|  | Omicron |  | N.E. |  |  |  |  | 2.1E-09 | 3.0E-10 | N.E. |

N. E. = no efficacy

Unfortunately, C5 showed no neutralizing activity against SARS-CoV-2 pseudovirus as the IC_50_ was greater than 1000 ng/ml (Figure 4B), probably due to its low binding affinity to RBD (1.80×10^-8^ M) (Figure 2F). In addition, as the protein yield and SEC profile were poor, we sought to engineer C5 by phage display to further improve its affinity and hopefully biophysical properties as well.

### Affinity maturation of C5 nanobody by phage display

To increase the binding affinity of nanobody C5 to RBD, we applied a method called “soft randomization” [37] . All the three CDRs of C5 were randomized simultaneously by doped oligos using Kunkel method, resulting in a bias library (See Methods). After four rounds of phage panning, ten C5 variants that specifically bind to RBD were obtained and their three CDRs were multi-aligned (Figure 3A). The multi-alignment result clearly showed the variable residues and the conserved ones in three CDRs. Notably, the variable residues of CDR3 gathered at the end of the loop, i.e. residues 104, 106 and 107. This highly variable region might enable CDR3 evolve extra binding capacity other than the regions at the central part.

**Figure 3.**
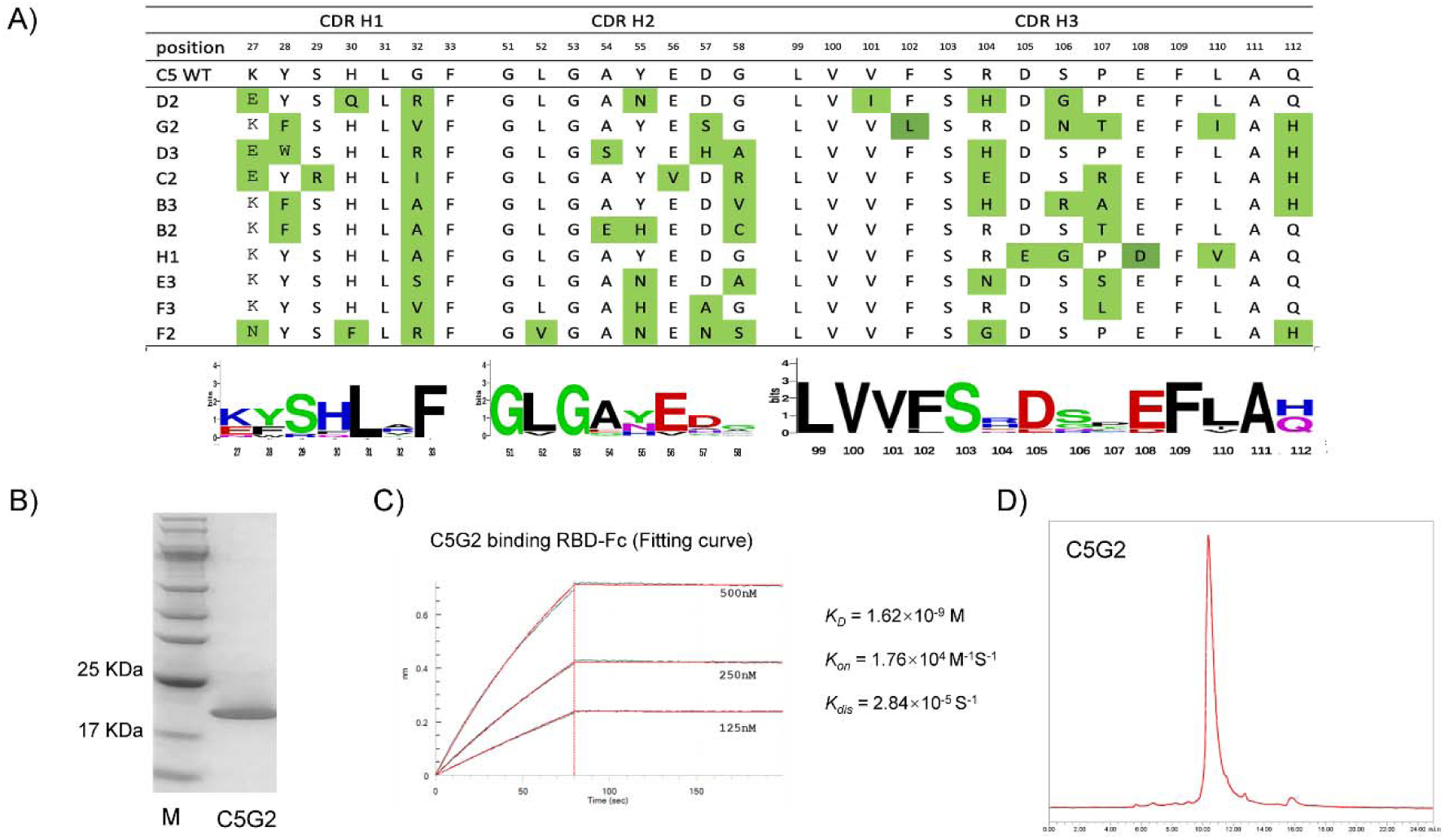
Affinity maturation of C5 clone by phage display led to the identification of C5G2 clone. A) C5 clone were used as template to generate an optimization library with mutation at all the CDRs. Panning of this library against RBD resulted in ten C5 variants. The CDR regions of these ten clones were aligned. The alignment represented by Weblogo were showed underneath each CDR region. B) C5G2 variant was purified from E. coli host and resolved in SDS-PAGE. C) Binding curve of C5G2 protein to RBD in BLI assay. The binding affinity increased from 18 nM to 1.62 nM. D) Size exclusion chromagraphy(SEC) profile of C5G2 protein resovled in a Tosho TSKgel G3000SW column.

As the C5 protein were highly insoluble (data not shown), we expected high yield of protein from those ten C5 variants. Fortunately, we were able to purify variant G2 (hereafter referred to C5G2) with significant increase in yield, i.e. from 0.3mg to 5mg protein per liter culture. The purified C5G2 was above 95% purity as resolved in SDS-PAGE (Figure 3B). The binding affinity of C5G2 to RBD (1.62×10^-9^ M) is ∼10 fold higher than the original C5 (Figure 3C), mainly due to the decreased K*_dis_* rate, which is from 2.59×10^-4^ s^-1^ to 2.84×10^-5^ s^-1^ (Table 2). The single peak in the SEC profile of C5G2 also indicated that there was no aggregation in this C5 optimized variant (Figure 3D).

### C5G2 showed broad-spectrum neutralizing activity against SARS-CoV-2 variants in pseudovirus assay

We then tested if the affinity matured C5G2 could inhibit the binding of RBD to ACE2 and subsequently neutralizing the SARS-CoV-2 pseudovirus infection. We also included D3 nanobody in this study as the negative control as well as previously reported nanobody mNb6 [10] as the positive control. We first applied a competitive ELISA assay, in which C5G2 protein and the control proteins were employed to compete with ACE2 binding to RBD. As shown in Figure 4A, with the increase of C5G2 protein concentration, the binding of RBD to ACE2 decreased and the measured IC_50_ was 3.75nM, which was comparable to that of mNb6 (4.64 nM). Meanwhile, the controlled D3 protein, which recognizes a non-RBD epitope, showed no inhibition at all. Based on the data, C5G2 nanobody could interfere with the binding of RBD to ACE2 and the efficient was closed to the reported potent nanobody mNb6.

**Figure 4.**
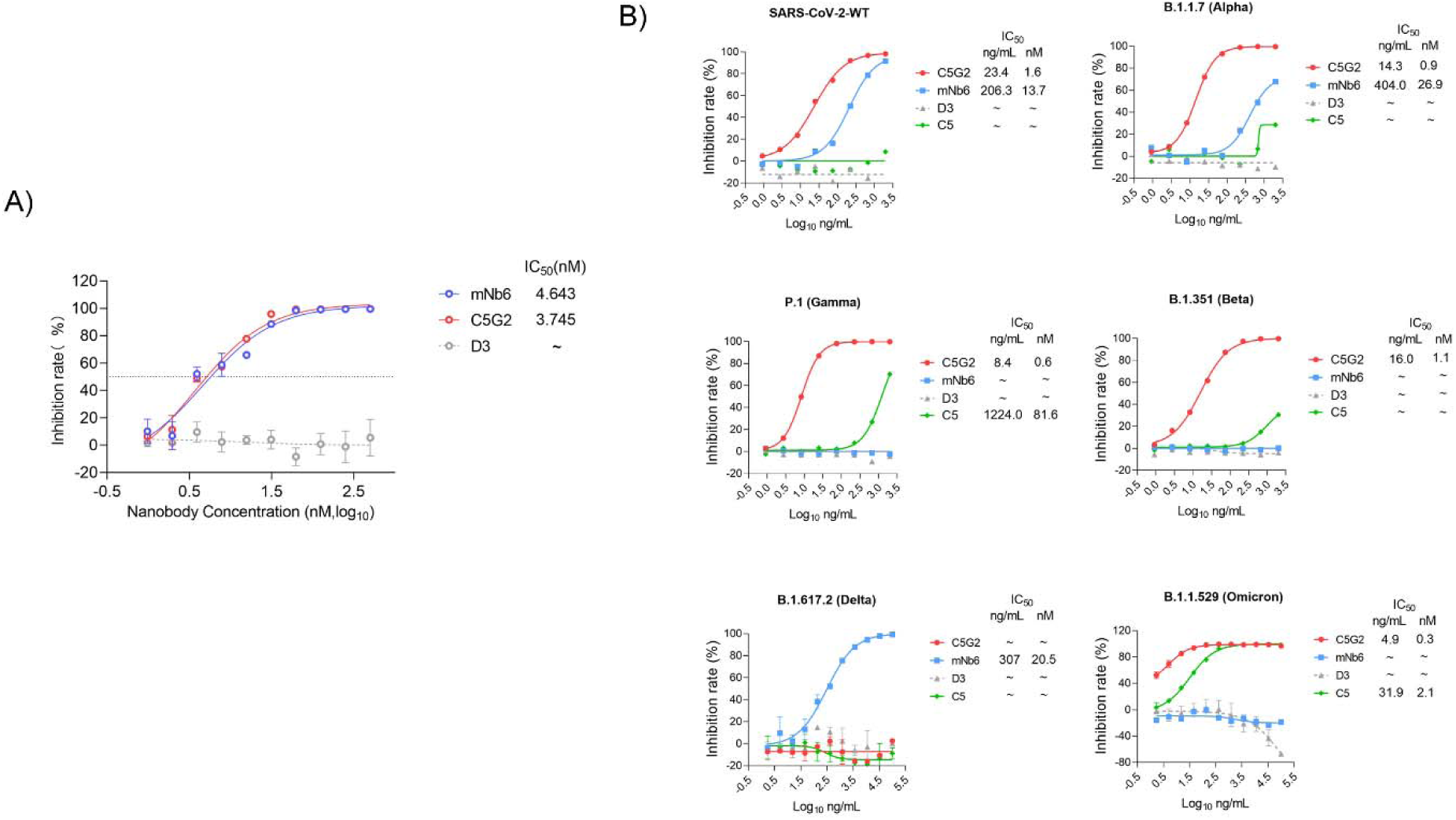
Pseudovirus assay of C5G2 nanobody. A) Competitive ELISA showed C5G2 protein, but not D3, competed with ACE2 to bind RBD with a IC50 of 3.75 nM. As a control, the IC50 of mNb6 for competition was 4.64 nM. The experiment was performed in tripilicated. B) Pseudovirus assay of C5G2’s inhibition potency to protect cells from virus infection. The wild type virus and Alpha, Beta Gamma, Delta and Omicron virus were test. mNb6 was a positive control. Nanobody D3 binding to NTD of S1 was the negative control. The IC_50_ value were showed both in ng/ml and nM(nmol/L).

We then applied pseudovirus assay to test if C5G2 protein could inhibit the infection of pseudovirus to living cells. As shown in Figure 4B, C5G2 could neutralize the WT pseudovirus with a high potency as the IC_50_ is 1.56nM. Meanwhile, the IC_50_ of mNb6 is 13.7 nM. C5G2 is about 9-fold higher potency than mNb6 in this experiment. Surprisingly, C5G2 showed even high potency in neutralizing Alpha, Beta and Gamma variants, with IC_50_ of 0.95 nM, 1.06 nM and 0.56 nM, respectively (Figure 4B). Unfortunately, C5G2 showed no neutralizing efficacy against Delta variant. During the preparation of this manuscript, the Omicron variant emerged and propagate rapidly worldwide. We produced the pseudovirus of Omicron variant and test C5G2’s potency in protecting the cells from virus infection. As showed in Figure 4B, C5G2 has an even higher potency as the IC50 of 0.3 nM compared to other VOCs. Taken together, C5G2 showed broad-spectrum neutralizing activities against most of VOCs except for Delta variant.

### Structural basis for the broad-spectrum neutralization activity of C5G2

To decipher the molecular basis for the neutralization breadth of C5G2, we determined the cryo-electron microscopy (cryo-EM) structure of C5G2 in complex with SARS-CoV-2 S protein at an overall resolution of 3.57 Å (Figure 5A-B, Figure S3 and Table S2). Compared to the wildly reported structure of prefusion spike which have one RBD in open (up) state and the other two in closed (down) state, C5G2 could recognize both the opened and the closed RBDs (Figure 5B). Due to its relatively smaller size than conventional IgG antibody (or Fab) and its long CDR3 loop, C5G2 could insert itself into the slit between the RBD and the adjacent NTD and make contact to both the RBD and NTD (Figure 5B). Unfortunately, because of the structural flexibility of the RBD, the interaction details were failed to unveiled in the initial reconstruction. Subsequently, we further performed a local refinement of one C5G2 plus the surrounding RBD and NTD (G5G2: RBD: NTD) at a resolution of 3.88 Å (Figure 5C, Figure S3 and Table S2). Consequently, the high resolution details of local refinement allowed us to build the atomic model of C5G2 and bound RBD and NTD, which confirmed the simultaneous interaction of C5G2 to the RBD and NTD (Figure 5D). C5G2 dominantly interacts with the RBD and results in an extremely large buried area of about 1,160 Å^2^ on RBD. Meanwhile, the buried area of the same C5G2 on the NTD is about 200 Å^2^ (Figure 5D).

**Figure 5.**
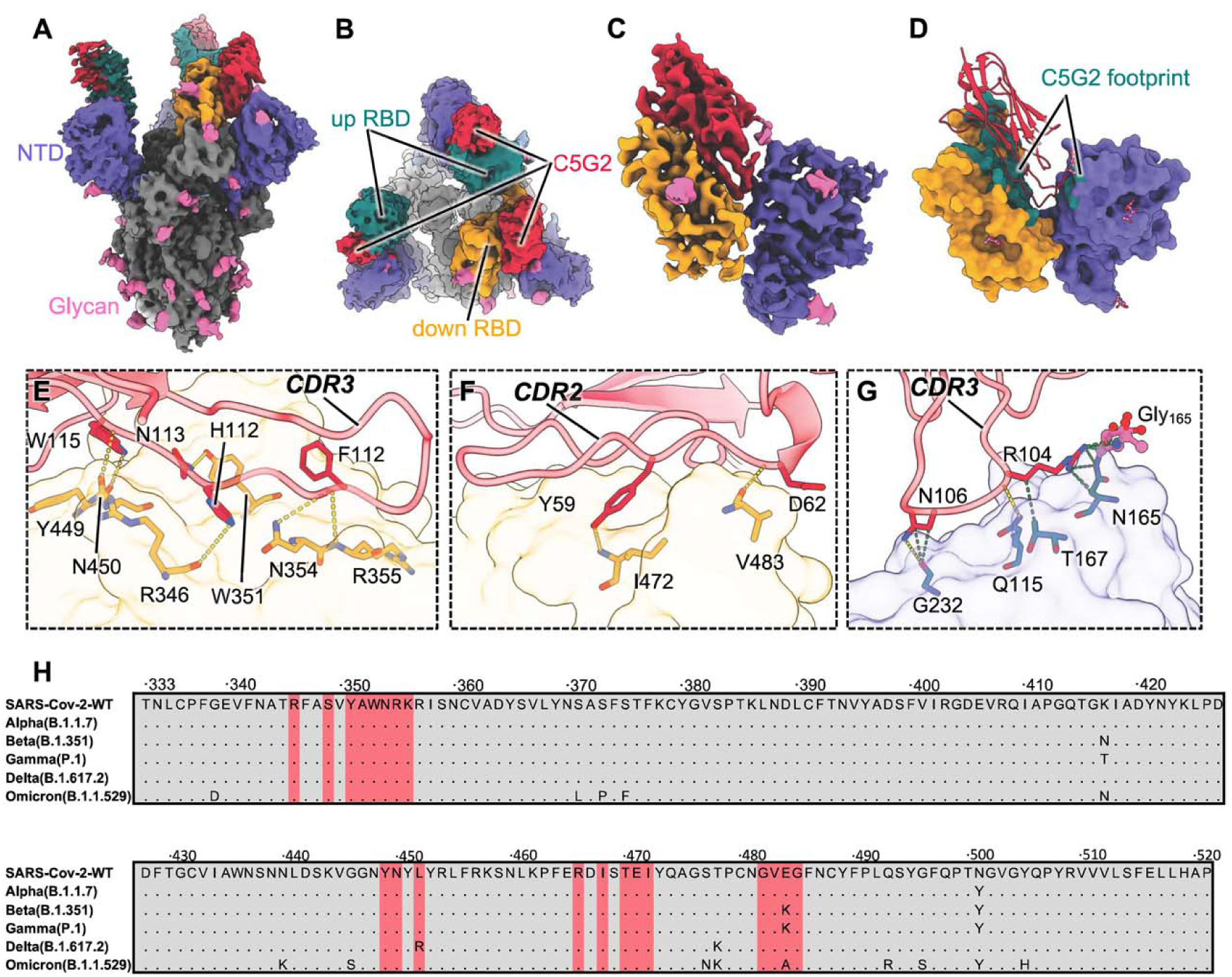
Structural basis for the broad neutralization of C5G2. **(A-B)** Cryo-EM density maps of SARS-CoV-2 spike protein in complex with C5G2 viewed from side (A) and top (B), respectively. C5G2 is colored in red and the up RBD, down RBD, NTD, glycans of spike are colored in teal, orange, blue and pink, respectively, the rest density of spike is colored in gray. **(C)** The 3.88 Å density map of local refinement of C5G2 and bound RBD and NTD. **(D)** Atomic model of G5G2:RBD:NTD. C5G2 is presented as cartoon and RBD and NTD are presented as surface representations, respectively. The epitopes of C5G2 on the RBD and NTD are both colored in teal. The decorated glycans on on RBD and NTD are shown as pink sticks. **(E-G)** the interaction details between C5G2 and RBD (E-F), NTD (G). C5G2 is showed as transparent cartoon and the RBD and NTD are shown as transparent surface. The residues involved in the hydrogen bond (gold dotted lines) interaction are shown as stick. The contacts between C5G2 and NTD are also presented as green dotted lines. **(H)** RBD sequences of SARS-CoV-2 WT and its 5 VOC variants (Alpha, Beta, Gamma, Delta and Omicron) with highlighted footprint of C5G2 (colored in light red).

C5G2 mainly uses its long CDR3 meditating the interactions with both the RBD and the NTD (Figure 5E-G). First, the CDR3 residues F109, H112, N113 and W115 form 5 hydrogen bonds to the RBD (Figure 5E). In addition, the CDR2 of C5G2 also interacts with the RBD and forms additional two hydrogen bonds mediated by the residues Y59 and D62 (Figure 5F). The framework 2(FR2) of C5G2 thus adopts an orientation that induce steric hindrance to the putative ACE2 that bind to the same RBD (Figure S4). As a result, C5G2 showed strong inhibition potential against the binding of ACE2 to RBD (Figure 4A). Furthermore, because of the long CDR3 loop, it extends to the neighboring NTD and formed 2 hydrogen bonds through residues R104 and N106 (Figure 5G). Of note, the NTD residue N165 and decorated glycan both participate in the interaction to C5G2 (Figure 5G). As the glycosylation of N165 functions as a switch of the up-to-down of RBD conformation[38], this interaction may interfere with the function of N165 glycan and enhance the inhibition of RBD binding to ACE2 subsequently.

Since it is the RBD that mainly mediated the interaction with C5G2, we next explored the conservation of the C5G2 epitope on RBD. A total of 20 residues (346, 349, 351-356, 449-450, 452, 466, 468, 470-472 and 482-485) on RBD are involved in the interactions to C5G2 (Figure 5H). Of these, 90% (18 residues) are highly conserved among all five variants of concern (VOCs) of Alpha, Beta, Gamma, Delta and Omicron (Figure 5H). Although the epitope residue L452 did not involved in hydrogen bond or salt bridge interactions to C5G2, the L452R mutation on Delta variant completely abolished the activity of C5G2 (Figure 4B). Furthermore, although the currently dominant Omicron variant holds unprecedented 15 mutations on its RBD, C5G2 showed effective neutralization potency against Omicron pseudovirus (Figure 4B). Structural analysis showed that, of this 15 mutations, only one (E484A) was found located in the footprint of C5G2 (Figure 5H), suggesting the epitope conservation among SARS-CoV-2 and Omicron variant.

## Discussion

Synthetic antibody library is an common and reliable way to obtain antibodies for diagnosis and therapy[10, 28, 39, 40]. In this study, we firstly constructed a universal synthetic nanobody library for the purpose of generating nanobodies for diverse antigens. The library design strategy is straightforward and readily be conducted. Though less sophisticated compared to the reported synthetic libraries [41, 42], the resulting library ASyNAL enabled us to obtain at least one nanobody for the randomly selected antigens (Table 1), proving its versatile utilization for nanobody generation. The binding affinity of the S protein (and its fragments) nanobodies from the ASyNAL library ranged from 5.58×10^-10^ M to 3.05×10^-7^ M (Table 2), which is within the affinity scope for the synthetic antibodies as well as the natural ones. The SEC profile of the S protein nanobodies indicated that most of the nanobodies have minimal aggregation. Therefore, the the ASyNAL library is a reliable resource of nanobodies with high affinities and good biophysical properties. Furthermore, the NGS data from library QC and panning results may help to design more sophisticated libraries for both naïve and engineered ones (e.g. affinity maturation).

We were able to obtained nanobody C5G2 against RBD of SARS-CoV-2 spike protein by panning the ASyNAL library and subsequent antibody engineering. C5G2 binds to S protein mainly through its CDR2 and CDR3 on a very conserved surface RBD in both up and down states(Figure 5B, H). In addition, the long CDR3 extends itself down to the bottom of the slit formed by RBD and neighboring NTD and makes contact with residues in NTD consequently (Figure 5D, G). This interaction may interfere with the function of N165 glycan and may further lock the RBD in the down state. Furthermore, the FR2 of C5G2 stick upward to RBM (receptor binding motif) to cause steric hindrance to prevent the potential ACE2 binding. This unique mechanism of C5G2 interaction with S protein underlies its basis of high potency of neutralization and broad activities(Figure 4A&B).

According to Hastie et al, the RBD antibodies could be classified into three group, those bind to RBM, outside face and inner face[43]. The RBM binders, usually compete with ACE2 to bind to RBD, lose(or weaken) their bindings and neutralizing potencies to the variants of SARS-CoV-2 virus, especially the most recent Omicron[44]. For example, the ultrapotent nanobody mNb6 lost its activities to most of the VOCs (Figure 4B) mainly due to its RBM binding property (Figure S5). In contrast, the clinically approved Vir-S309 monoclonal antibody still neutralized Omicron with a slightly potency decrease (VSV IC_50_=284ng/mL) as it binds to a conserved and outer face of RBD (Figure S5). The C5G2 nanobody in this study binds to a RBD epitope with >90% residue conserved in the VOCs and 80% residues conserved in all the SARS-CoV-2 variants reported so far(Table S3), underlying its broad-spectrum activity against most of the VOCs (Figure 4B). Moreover, we found C5G2 nanobody could protect cells from swine coronavirus infection in a separate experiment(data not shown), implying its epitope conservation and broad activity across coronavirus species.

N165 is a conserved residue at NTD and its glycan plays a pivotal role in facilitate the transition of RBD from down to up state conformation and stabilize the conformation by interacting with RBD residues Y351, T470, F490, and L452 [38, 45]. C5G2 can not only interact with N165 glycan in NTD (Figure 5G), but also three of the four epitopic residues (i.e. Y351, T470, and L452) for N165 glycan in RBD (Figure 5H). This interaction of C5G2 could block N165 glycan binding to RBD as exemplified by neutralizing antibody 35B5[45]and stabilize RBD in down state, thus enhance the the inhibition efficiency of C5G2. The ACE2 interacts RBD in up state during virus infection. However, the steric hindrance caused by FR2 of C5G2 upon binding could further inhibit the ACE2 binding to RBD. So C5G2 actually acts as a triple-function nanobody, i.e. RBD binder, NTD binder and ACE2 blocker(Figure S4). As a result, monomeric C5G2 nanobody inhibited most of the VOCs with high potency as the IC_50_ from 4.9-23 ng/mL in pseduovirus assay (Figure 4B). Furthermore, the potency of C5G2 could be driven even higher by linking two or three nanobodies together, e.g. the case of bi-mNb6 and tri-mNb6[10].

Unfortunately, the advantage of C5G2 for virus inhibition was compromised by the L452R mutation of RBD in the variant of Delta (Figure 4B). Linking C5G2 with a heterogeneous nanobody may compensate for this disadvantage as exemplified by the bi-specific sdAb bn03(Figure S5)[46]. Due to its renewable characteristics, further engineering of the CDR loops of C5G2 by directed evolution may also find a new nanobody variant to tolerate this mutation while keeps the other functions. Taken together, this monomeric, small size(15kD) nanobody can compete with full IgG and bi-specific nanobody(Figure S5, lower table) in preventing current SARS-CoV-2 and its variants through its unique mechanism of action. Due to the highly conserved epitopes in RBD and NTD, C5G2 also holds the promise to inhibit the emerging virus in the future.

## Materials and Methods

### Construction of a synthetic nanobody library

In order to construct a novel synthetic nanobody library, a natural nanobody (PDB no. 1ZVH) from camel that recognizes lysozyme was used as template for mutagenesis. The coding cDNA of 1ZVH was chemically synthesized and subcloned into a modified phage display vector of pComb3XSS, in which the amber stop codon (TAG) were mutated to CAG to facilitate nanobody display in *E. coli* without gene supE, e.g. SS320(Genentech). The 1ZVH construct was firstly transformed into *E. coli* SS320 and the phage were produced by adding helper phage M13KO7 (New England Biolabs). The successful display of 1ZVH was then verified by phage ELISA, in which the produced phage could be bound by immobilized lysozyme in a 96-well NUNC Maxisorp plate.

To increase the oligo incorporation efficiency, two ochre stop codon (TAA) and one BamH I restriction enzyme site (GGATCC) in between were applied to replace the CDR3 region of 1ZVH to generate a so called “Stop template”. The “Stop template” was transformed into *E. coli* CJ236 (New England Biolabs) to produce *du* single strand DNA (*du*-ssDNA) as described previously[47]. The mutation strategy was using degenerate oligo mix (X) to incorporate into the CDR1 (7aa), CDR2 (8aa) and CDR3 (14aa) of the 1ZVH, respectively (Figure 1A). The degenerate oligo mix X denotes 16% of Tyrosine, 10% of Glycing, 10% Serine, 5% of Alanine, Aspartate, Glutamate, Phenylalanine, Histidine, Isoleucine, Lysine, Leucine, Glutamine, Arginine, Threonine, and Valine, respectively and 1% of Methionine, Asparagine and Tryptophan, respectively. The oligos for each CDR were synthesized by a TRIMER-like method by Genewiz (Suzhou, China). The CDR oligos were incorporated into the corresponding CDR regions simultaneously by standard Kunkel reaction as described previously[36]. The Kunkel product was transformed into 350 µL high efficiency competent cell SS320 (with M13KO7 incorporated) made by electroporation as described previously [36]. The transformed SS320 were grown overnight in 2YT media supplemented with 50 μg/mL carbenicillin at 32°C with a shaking speed of 200 rpm. To increase the incorporation percentage at CDR3 region, the SS320 cells were pelleted and the the plasmid DNA were maxipreped. 20 µg plasmid DNA were digested by restriction enzyme BamH I to remove those DNA molecules with failed incorporation at CDR3 region (i.e. the Stop templates) for at least 4 hours at 37°C. The resultant DNA were purified in a PCR product purification column to remove ions and enzymes. The DNA sample were re-suspended into a minimum volume, i.e. 60µL H_2_O. The digested DNA sample was re-transformed into 350 µL competent cell SS320 (+M13KO7) by electroporation. The transformed SS320 cells were grown as above and the phages in the supernatant were precipitated by PEG/NaCl (20%/2.5M). The concentration of the phage solution was determined by serial dilution and by infection of XL1-Blue (Stratagene) as described previously[36].

### Library quality control by next generation sequencing

The library DNA were applied as templates in PCR reaction and the nanobody encoding sequences were amplified by Phanta Max Super-Fidelity DNA Polymerase (Vazyme). The PCR products were resolved in 1.5% agrose gel and the band was sliced for gel purification. The purified DNA were subject for Next generation sequencing (NGS). More than 50 ng purified PCR fragments were used for library preparation. These PCR products were treated with End Prep Enzyme Mix for end repairing, 5’ Phosphorylation and dA-tailing in one reaction, followed by a T-A ligation to add adaptors to both ends. Adaptor-ligated DNA was then performed using DNA Clean Beads. A second PCR reaction was carried out with two primers carrying sequences which can anneal with flowcell to perform bridge PCR and index allowing for multiplexing. The final library product for sequencing were then purified by beads and was qualified. The qualified libraries were sequenced pair end 150 bp on lllumina NovaSeq 6000. The method of data analysis is performed as described previously [36].

### Phage biopanning

The phage biopanning was performed as previously described [48]. Briefly, Two wells of 96-well microplate (NUNC) were coated with 0.5 μg antigen (e.g. RBD-Fc or other antigens) and 0.5μg Fc in 100μL 1×PBS (137 mM NaCl, 3 mM KCl, 8mM Na_2_HPO_4_ and 1.5mM KH_2_PO_4_, pH=7.6) overnight, respectively. Solution in the well were discarded and 200 μL/well 2% skim milk were added for blocking at room temperature for 1 hour. After incubation, solution were discarded and 100 μL/well phage-displayed nanobody library (∼3.0×10^12^ phage clones, approximately 1000× library diversity) were added to the Fc wells for pre-incubation to remove the non-specific binders. After incubation at room temperature for 1 hour, phage solution were then transferred into the RBD-Fc well for binding for 1 hour. Non-binding phages were washed away by PT buffer (1×PBS + 0.05% Tween) for at least 8 times. Bound phages were eluted by 100mM HCl 100 μL/well. 1/8 volume of Tris-HCl (1M, pH=11) were then added for neutralization. Half volume of the neutralized phage solution was then applied to infect 10-fold volume of actively growing *E. coli* XL1-blue (Stratagene) for 30 min at 37 °C. Then M13KO7 helper phages (NEB, N0315S) were added at a final concentration of 1×10^10^ phages/mL for super infection for 45 min. The XL1-blue culture was added into 20-fold volume of 2YT medium (10 g yeast extract, 16 g tryptone, 5 g NaCl in 1L water) supplemented with Carb (carbenicillin, 50 mg/μL) and Kana (kanamycin, 25 mg/μL), and grow overnight (14-16 hours) at 32°C, 200 rpm in a shaker. The overnight culture was centrifuged and the supernatant were precipitated by 1/5 volume of PEG/NaCl (20% PEG 8000/2.5M NaCl). The amplified phage in the pellets were resuspend with 1mL 1×PBS and applied as the input phage for the next round of panning. For C5 variant library panning, from the second round, antigen RBD-Fc were decreased from 0.5μg to 0.2μg (2^nd^ round), 0.1μg (3^rd^ round), 0.05μg (4^th^ round) to increase the selection stringency.

### Phage ELISA

In a 96-well NUNC microplate, 0.1 μg /well RBD-Fc (or other antigens) and 0.1 μg /well Fc were coated in wells in 50μL 1×PBS, respectively. After incubation at 4°C overnight, solution in the well was discarded and 100 μL/well 2% milk were added for blocking at room temperature for 1 hour. 2×50μL phage solution were added into the RBD-Fc wells and Fc wells, respectively, for binding for 1 hour. Non-binding phage were washed away 8 times by PT buffer. 50μL anti-M13/HRP conjugate (Sino Biological) were added and incubated for 30 min. After wash by PT buffer, 50μL TMB substrate were add to develop color according to the manufacturer’s instruction. 100 μL of 1.0 M H_3_PO_4_ were added to stop the reaction and signals were read spectrophotometrically at 450 nm in a BioTek plate reader. The reads of RBD-Fc and Fc wells were recorded and the ratio of RBD-Fc/ Fc were calculated.

### Competitive phage ELISA

0.2 ug/well Spike-trimer-His (Acrobiosystem) /RBD-Fc were coated in microplate in 50uL 1×PBS buffer at 4°C overnight. 100 μL/well 2% milk were incubated for 1 hour for blocking. After washing for 3 times, a serial phage solution was added to incubate with spike-trimer-His for 30 minutes. Anti-M13/HRP conjugate (1:8000 dilution) and TMB were used to amplify the signals and develop color reaction, respectively. After stopping reaction via 1.0 M H_3_PO_4_, OD 450 value was read by microplate reader. After that, subsaturation (80% of maximal effect) concentration of phage solution was used for competitive Elisa.

0.2 ug/well Spike-trimer-His/RBD-Fc were coated for competitive phage Elisa. After incubation with 100 uL/well 2% milk for 1 hour and 3 times washing, subsaturation concentration of phage solution mixed with decreased concentration of ACE2-Fc (Novoprotein). The mixture of phage and ACE2-Fc was added to wells coated with Spike-trimer-His/RBD-Fc and incubation for 30 minutes. M13/HRP conjugate, TMB, 1.0 M H_3_PO_4_ and microplate reader were used for subsequent operation as described above.

### C5 affinity maturation library construction

The sequence of C5 in phagemid pComb3XSS were used as template for library construction. The C5 du-ssDNA used for Kunkel reaction was made as described previously [47]. Primer 1 (AGCTGTGCAGCAAGTGGA*<u>TAAGGATCCTAA</u>*CTAGGCTGGTTTCGTCAA), Primer 2 (CGCGAAGGAGTTGCTGCA*<u>TAAGGATCCTAA</u>*TACTACGCCGATAGCGTG), and Primer 3 (CTGTACTATTGTGCGGCC*<u>TAAGGATCCTAA</u>*AACTACTGGGGCCAAGGC) were used in a combinatorial reaction by Kunkel method to construct a “Stop template”, in which all CDRs were incorporated with BamH I restriction enzyme recognition site *<u>GGATCC</u>* and two stop codons *<u>TAA</u>*. Then Primer 4 (AGCTGTGCAGCAAGTGGAN_1_N_1_N_1_N_2_N_1_N_2_N_1_N_4_N_3_N_3_N_1_N_2_N_3_N_2_N_4_N_4_N_4_N_2_N_2_N_2_N_2_CTAGGCTGGTTTCGTCAA), Primer 5 (CGCGAAGGAGTTGCTGCAN_4_N_4_N_2_N_3_N_2_N_4_N_4_N_4_N_2_N_4_N_3_N_1_N_2_N_1_N_2_N_4_N_1_N_1_N_4_N_1_N_2_N_4_N_4_N_2_TACTACGCCGATAGCGTG) and Primer 6 (CTGTACTATTGTGCGGCCN_3_N_2_N_4_N_4_N_2_N_2_N_4_N_2_N_2_N_2_N_2_N_2_N_1_N_4_N_3_N_3_N_4_N_2_N_4_N_1_N_2_N_1_N_4_N_3_N_3_N_3_N_4_N_4_N_1_N_1_N_2_N_2_N_2_N_3_N_2_N_4_N_4_N_3_N_1_N_3_N_1_N_4_AACTACTGGGGCCAAGGC) were applied simultaneously in a Kunkel method to synthesize the heteroduplex double strand DNA (dsDNA), in which the designed mutations at each position were incorporate into the “Stop template”. In the above primers, N_1_ is composed of the mixture of 85% of A, 5% of G, T and C, respectively; N_2_ represents the mixture of 85% of T and 5% of A, G and C; N_3_ means the mixture of 85% of C, 5% of A, T and G; N_4_ contains 85% of G, 5% of T, A and C. The dsDNA was further digested by restriction enzymes BamH I to remove the unreacted template molecules before transformed into *E. coli* SS320 (pre-infected by M13KO7) by electroporation. The transformation efficiency (library diversity) was calculated by bacterial serial dilution as described [48]. The resulting phage library was precipitated by PEG/NaCl (20% PEG 8000/2.5M NaCl) as described[36].

### Nanobody expression and purification

The cDNA encoding the nanobodies in the pComb3XSS vector were PCR amplified and subcloned into vector pET22b to express 6×His tagged proteins. The expression constructs were transformed in to *E. coli* BL21 (DE3). Single colonies were picked and grow in 2YT/Carb medium at 37°C to OD_600_=0.8. IPTG were added to a final concentration of 0.2 mM and protein expression was induced at 18 °C overnight. The culture was centrifuged and the pellets were treated as previously described [49]. Briefly, pellets were re-suspended in lysis buffer (to make 100 mL lysis buffer: 98 mL Hepes/NaCl (50 mM/500mM) buffer, 1 mL TritonX-100, 1 mL 100× protease inhibitor cocktail, 10 μL benzonase, 5% glycerol, 100 mg lysozyme). Both the lysate and the supernatant from the overnight were heated at 60°C for 30 min to denature partially folded nanobodies. Heat-treated lysate and supernatant were centrifuged again and subjected for antibody purification. Antibody were purified using Ni-NTA agrose (Qiagen) according to the manufacturer’s manual. The eluted proteins were buffer exchanged into 1×PBS by Amicon Ultra-4 Centrifugal Filter Units (Millipore). The final concentrations of the proteins were determined by BCA method. The purity of the nanobodies were resolved in 15% SDS-PAGE gels.

### Competitive protein ELISA

In a 96-well NUNC microplate, 0.2 μg ACE2-His (Novoprotein) were coated per well in 50 μL 1×PBS at 4°C overnight. Then 100 μL/well 2% milk were added for blocking for 1 hour. A serial RBD-Fc with increased concentration (0nM, 3.125nM, 6.25nM…100nM, 200nM) were added in 9 wells coated with ACE2-His. The wells were washed 5 times by PT buffer after incubation for 1 hour at room temperature. 50 μL of Anti-IgG-HRP conjugate (Shanghai Ruian) were added to each well and incubated for 30 min. After washing 8 times by PT buffer, 50μL TMB substrate were add to develop color for 2 min. 100 μL of 1.0 M H_3_PO_4_ were added to stop the reaction and signals were read spectrophotometrically at 450 nm in a plate reader. Standard variation values were calculated using a 3-parameter logistic regression fit using Prism Software (GraphPad). Concentration for 80% of maximal effect (the sub-saturation concentration) were applied for the following competitive ELISA. 0.2μg/well ACE2-His were coated in a 96-well NUNC microplate. After incubation at 4 °C overnight, 100 μL/well 2% milk were used for blocking. Sub-saturation concentration of RBD-Fc were mixed with a serial nanobody-His with decreased concentration (500nM, 250nM, 125nM… 0nM). The mixture was added to 12 wells coated with ACE2-His and incubated at room temperature for 30 mins. After incubation with Anti-IgG-HRP and washed by PT buffer, TMB were used to develop color. The reaction was terminated with 1.0 M H_3_PO_4_ and the OD_450_ value was read through a Biotek Microplate Reader. Each reaction was triplicated and the mean of the readout were used for IC_50_ calculation.

### Size-exclusion chromatography

The buffer of nanobody was changed to 1×PBS (PH=7.4) by ultrafiltration discs (Merck, PLBC07610). 50μL nanobody (1 mg/mL) were filtered through a 0.2 μm filter. After that, filtered samples were prepared for HPLC analysis. HPLC instrument (Waters, ACQUITY UPLC H-Clas) was used to purity analysis of proteins. PBS and the gel permeation chromatography column (TSKgel G3000SW_XL_,Tosoh Bioscience, 0008541) were used as the mobile phase and the stationary phase, respectively. The pump was running for 5 minutes, mainly to eliminate air bubbles in the system, and then closed all exhaust valves. The mobile phase was running at a fixed rate(1mL/min) until the baseline is stable. Then, the operating parameters(Flow rate: 1 mL/min, pressure: 3.3MPa, temperature: 25°C, detection: UV 280nm) of sample was set in the software (Waters, Empower3). After that, sample was injected into the sampling valve and the line was monitored by software. The sample was allowed to run for 20 min (more than 1.5✕column volume ) before stopped.

### Bio-layer interferometry (BLI ) assay

BLI experiments were carried out using an Octet RED96 System (ForteBio). The measurements were performed using Ni-NTA biosensors (ForteBio). The nanobodies with 6×His tag were immobilized on the biosensor tip surface. All steps were performed at 30°C with shaking at 1000 rpm in a black 96-well plate, with a working volume of 200 μL in each well.

The RBD-Fc with different concentration (250nM, 125nM, 0nM) in the running buffer (1×PBS + 0.5% BSA + 0.05% Tween) was applied for association and dissociation. The response data were normalized using Octet data analysis software version 9.0.0.14 (ForteBio).

### Pseudovirus-based neutralization assay

Vesicular stomatitis virus (VSV) dG-SARS-Cov2 virus were used as pseudovirus and BHK21-hACE2 were prepared for neutralization assay [50]. Gradient diluted nanobodies were added to VSV dG-SARS-Cov2 virus (MOI=0.05) then incubated at 37°C for 1h. All samples and virus were diluted by 10% FBS-DMEM. After incubation, mixture were added to BHK21-hACE2 and incubated for 12h. Fluorescence images were obtained with Opera Phenix or Operetta CLS equipment (PerkinElmer) and analyzed by the Columbus system (PerkinElmer). Compare the number of GFP-positive cells in nanobody treated group and in nontreated group to calculate the reduction ratio which represents the neutralizing potency.

### Cryo-EM sample and data collection

Aliquots (3 μL) of 3.5 mg/mL mixtures of purified SARS-CoV-2 S protein (Acrobiosystem) in complex with excess C5G2 of three nAbs were incubated in 0.01% (v/v) Digitonin (Sigma) and then loaded onto glow-discharged (60 s at 20 mA) holey carbon Quantifoil grids (R1.2/1.3, 200 mesh, Quantifoil Micro Tools) using a Vitrobot Mark IV (ThermoFisher Scientific) at 100% humidity and 4°C. Data were acquired using the SerialEM software on an FEI Tecnai F30 transmission electron microscope (ThermoFisher Scientific) operated at 300 kV and equipped with a Gatan K3 direct detector. Images were recorded in the 36-frame movie mode at a nominal 39,000× magnification at super-resolution mode with a pixel size of 0.389 Å. The total electron dose was set to 60 e^−^ Å^−2^ and the exposure time was 4.5 s.

### Image processing and 3D reconstruction

Drift and beam-induced motion correction were performed with MotionCor2 [51] to produce a micrograph from each movie. Contrast transfer function (CTF) fitting and phase-shift estimation were conducted with Gctf [52]. Micrographs with astigmatism, obvious drift, or contamination were discarded before reconstruction. The following reconstruction procedures were performed by using Cryosparc V3 [53]. In brief, particles were automatically picked by using the “Blob picker” or “Template picker”. Several rounds of reference-free 2D classifications were performed and the selected good particles were then subjected to ab-initio reconstruction, heterogeneous refinement and final non-uniform refinement. The resolution of all density maps was determined by the gold-standard Fourier shell correlation curve, with a cutoff of 0.143 [54]. Local map resolution was estimated with ResMap [55].

### Atomic model building, refinement, and 3D visualization

The initial model of nAbs was generated from homology modeling by Accelrys Discovery Studio software (available from: URL: https://www.3dsbiovia.com). The structure of SARS-CoV-2 RBD from the structure of WT trimeric spike (pdb no. 6VSB[56]) was used as the initial modes of our WT-RBD. We initially fitted the templates models into the corresponding final cryo-EM map using Chimera [57], and further corrected and adjusted them manually by real-space refinement in Coot [58]. The resulting models were then refined with phenix.real_space_refine in PHENIX [59]. These operations were executed iteratively until the problematic regions, Ramachandran outliers, and poor rotamers were either eliminated or moved to favored regions. The final atomic models were validated with Molprobity [60, 61]. All figures were generated with Chimera or ChimeraX [62, 63].

