## Supplemental Tables&Figures for "A potent synthetic nanobody with broad-spectrum activity neutralizes SARS-Cov-2 virus and Omicron variant through a unique binding mode"

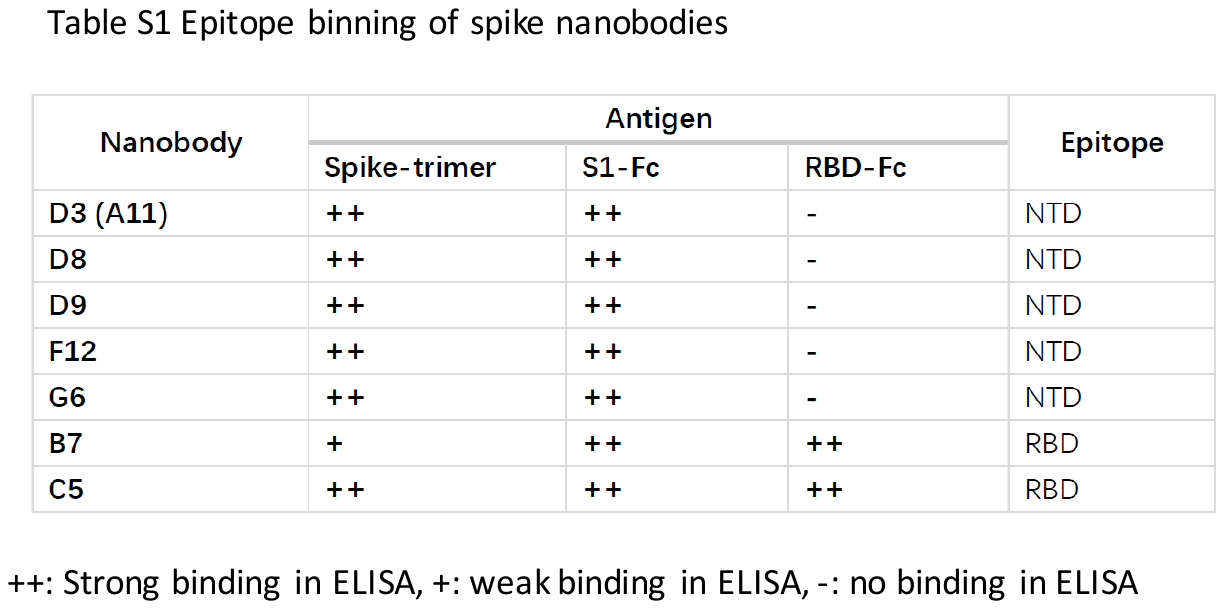

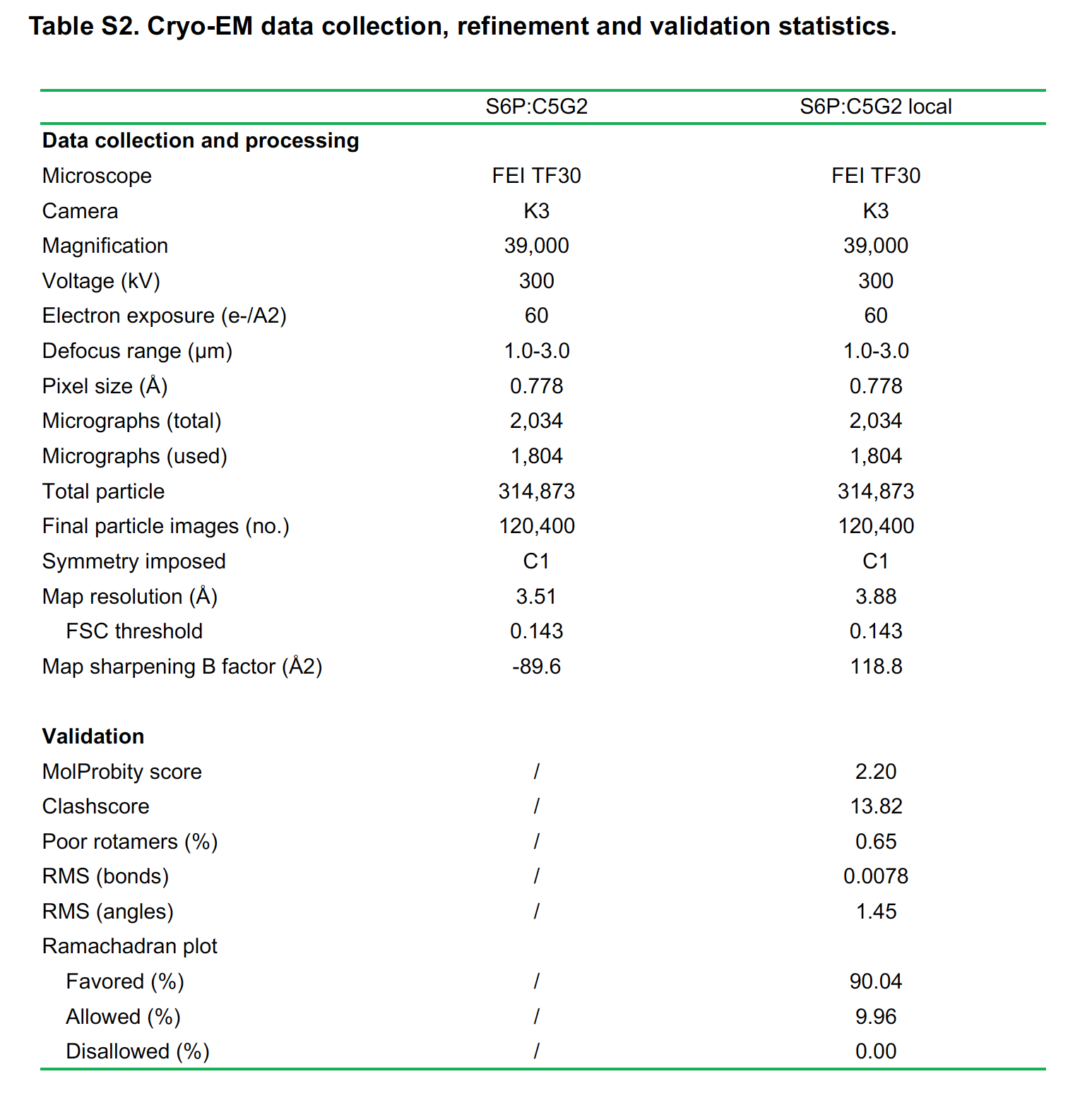

**Table S3 Conservation of epitopic residues in RBD for C5G2 binding from natural occurring SARS-Cov-2 variants**

| Epitopic residues of C5G2 in RBD | Occurency mutation(%)* |
| --- | --- |
| 346 | 1-10 |
| 349 | 0 |
| 351 | 0 |
| 352 | 0 |
| 353 | 0 |
| 354 | 0-0.1 |
| 355 | 0 |
| 356 | 0 |
| 449 | 0 |
| 450 | 0 |
| 452 | >10 |
| 466 | 0 |
| 468 | 0 |
| 470 | 0 |
| 471 | 0 |
| 472 | 0 |
| 482 | 0 |
| 483 | 0 |
| 484 | 1-10 |
| 485 | 0 |

* n=29624, data reproduced from Li et al [4].

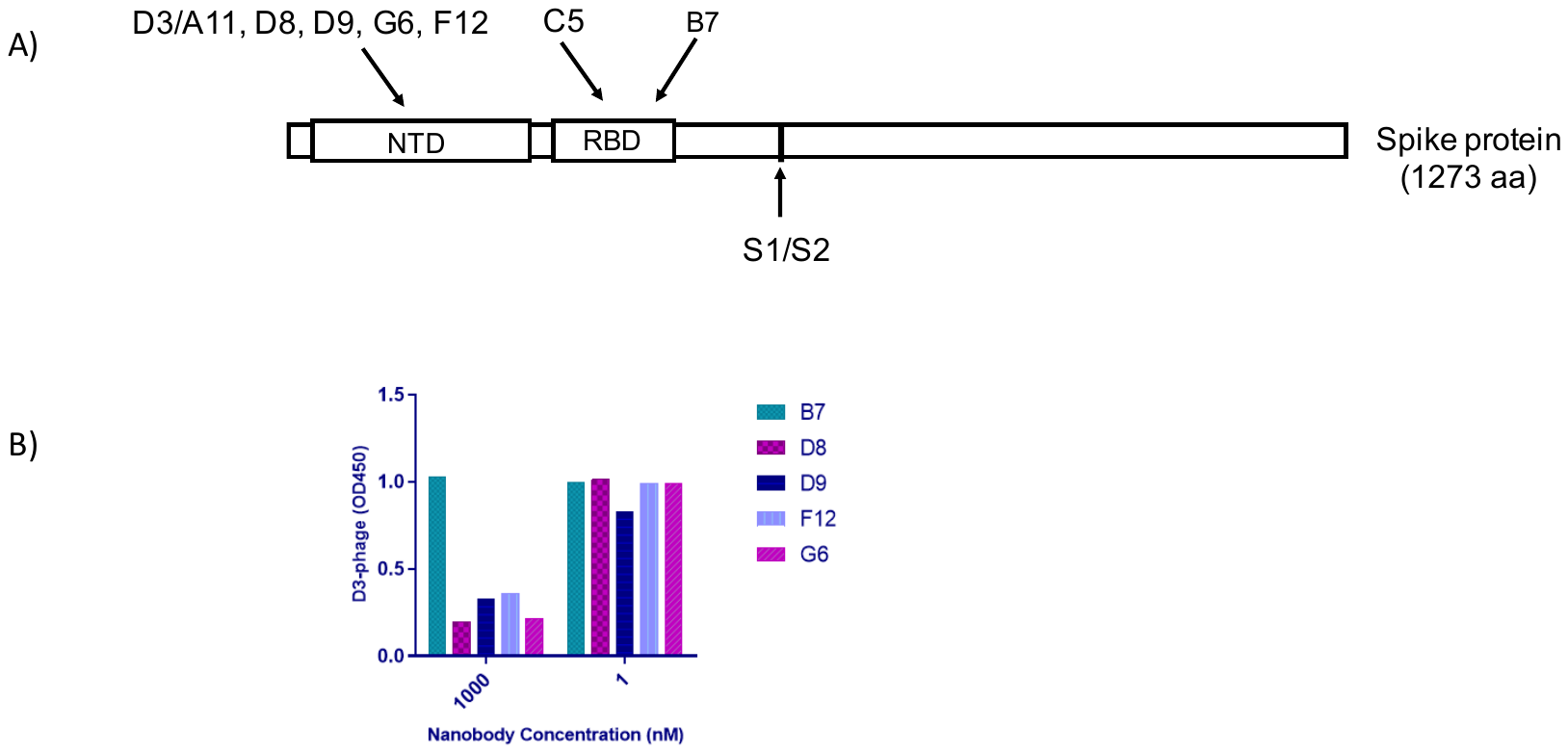

**Figure S1 Nanobodies selected by S/S1/RBD protein binding to different epitopes**

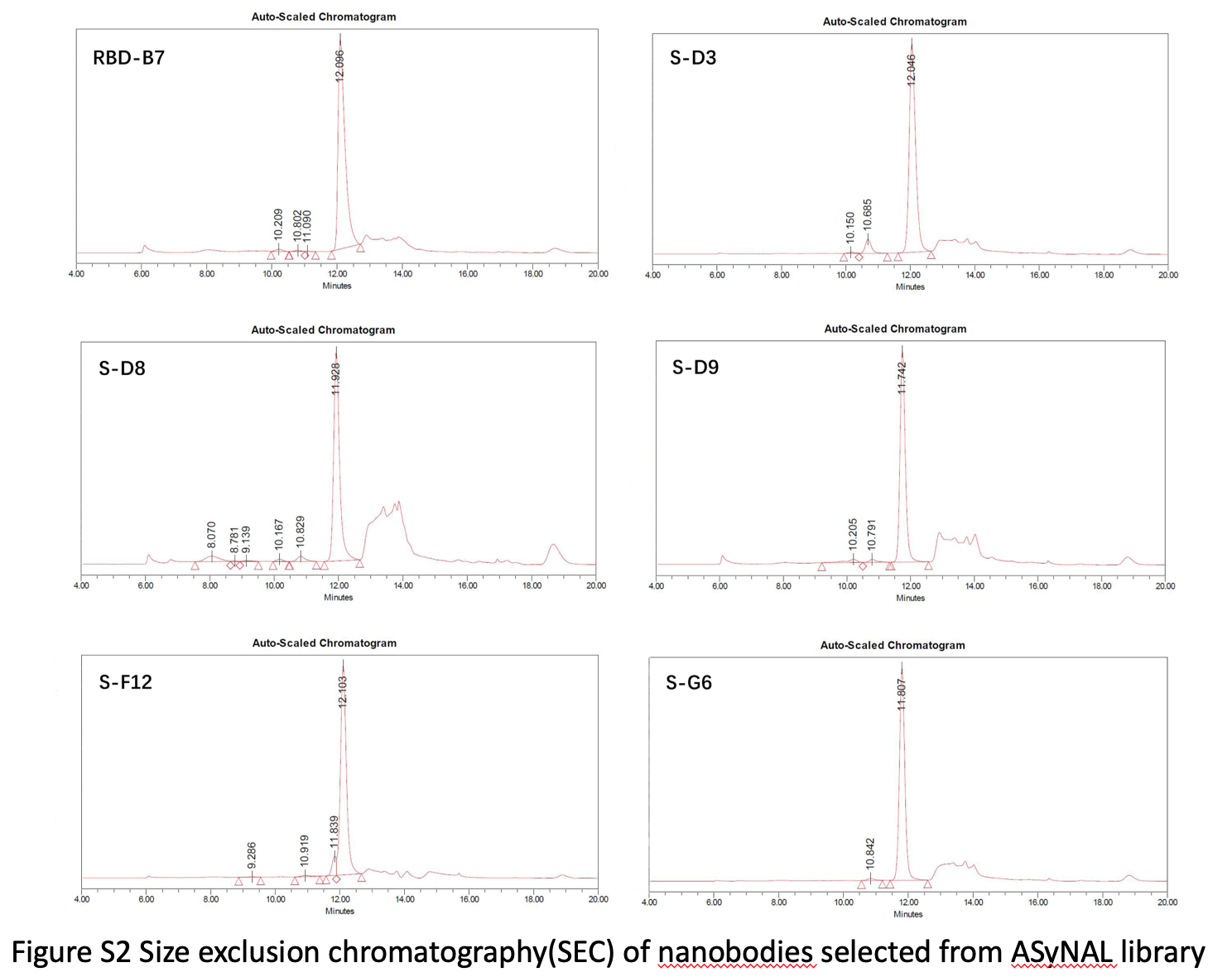

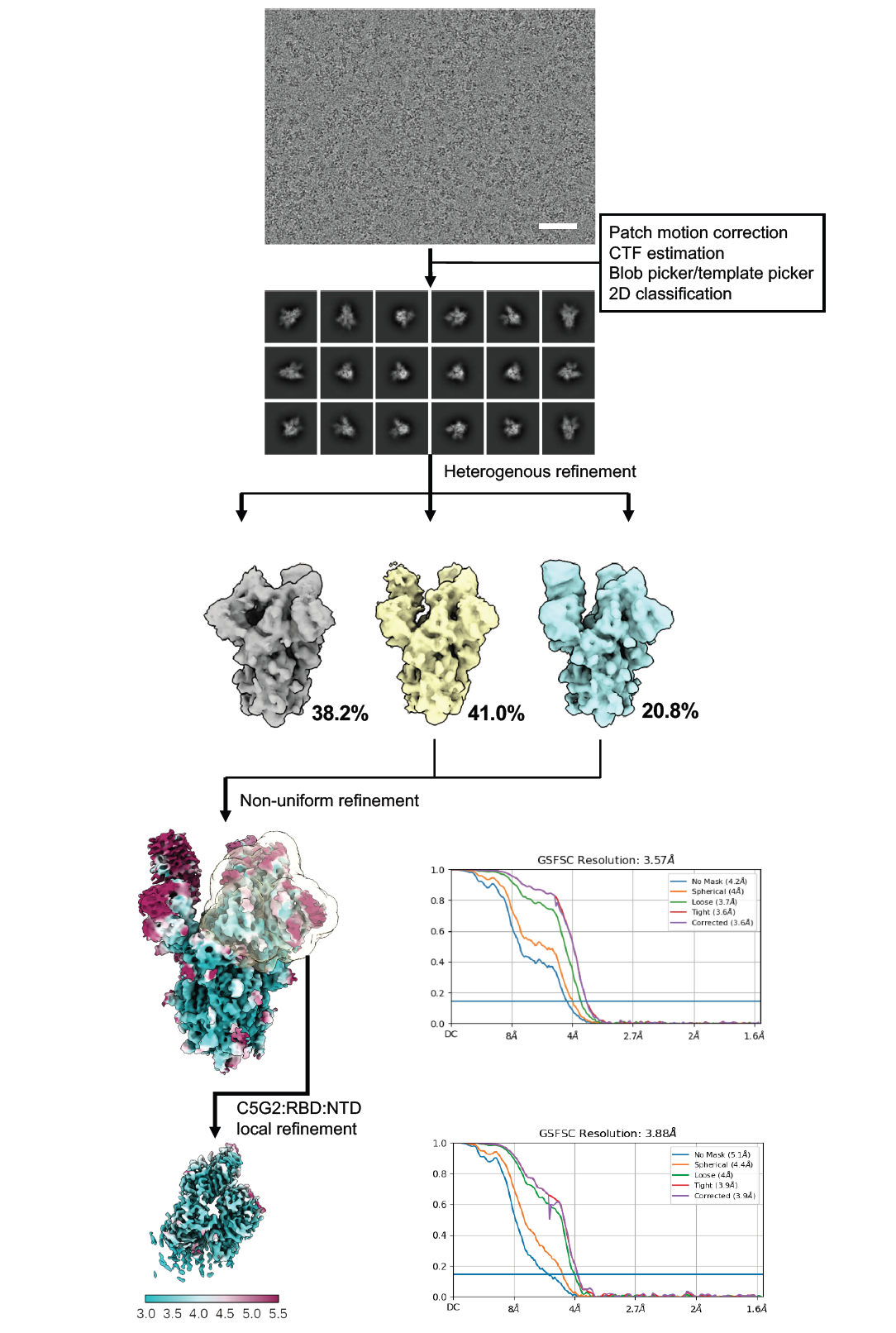

**Figure S3 Single-particle cryo-EM images processing workflow and the global and local resolution estimation for the immune complex of SARS-CoV-2 WT-S6P:G5G2.**

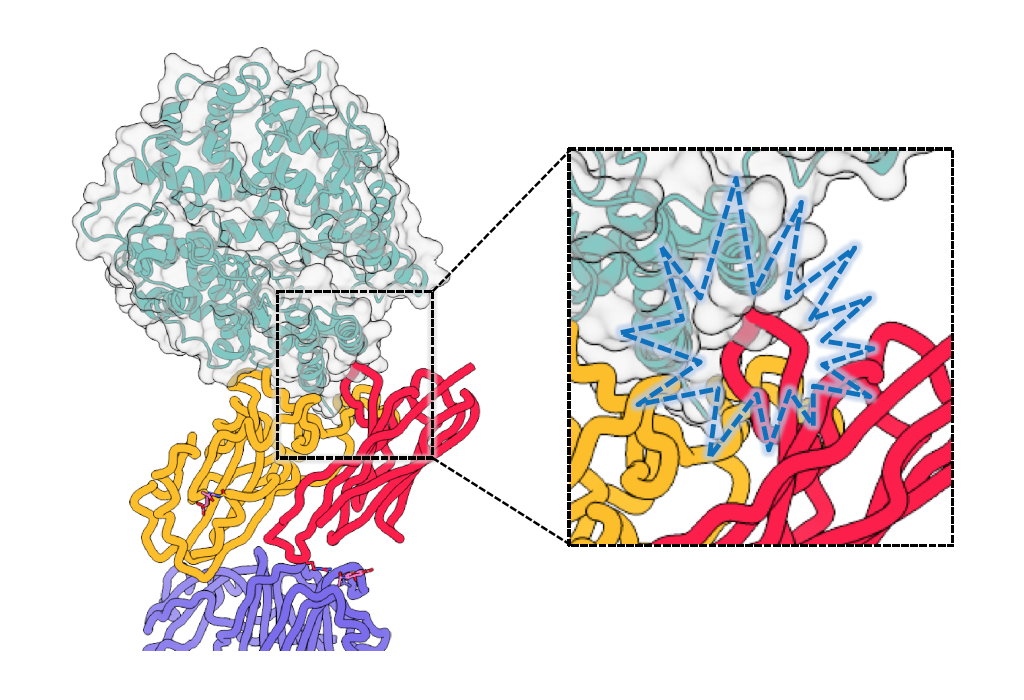

**Figure S4 Potential steric hindrance between C5G2 and ACE2 when binding to RBD simultaneously.** The RBD, C5G2, ACE2 and NTD were colored in yellow, red, turquoise

and blue, respectively.

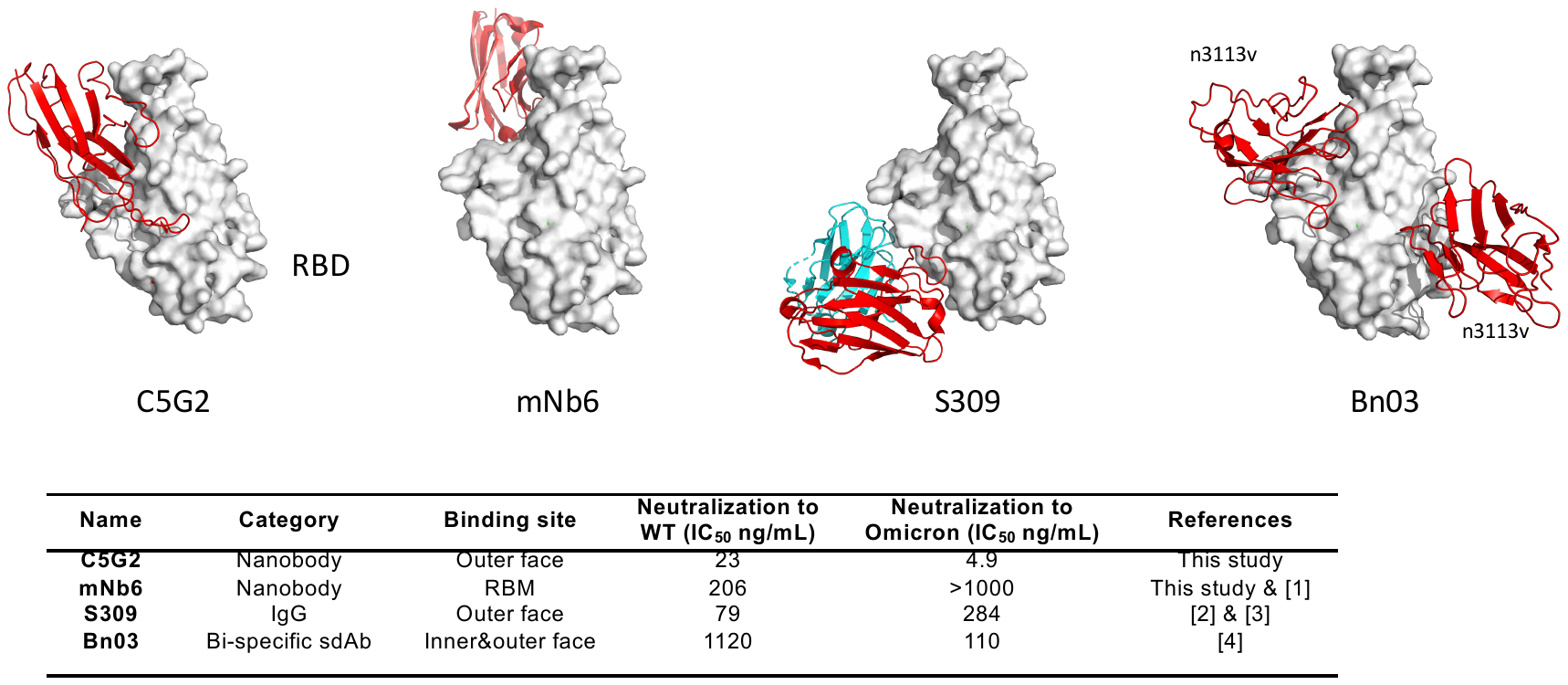

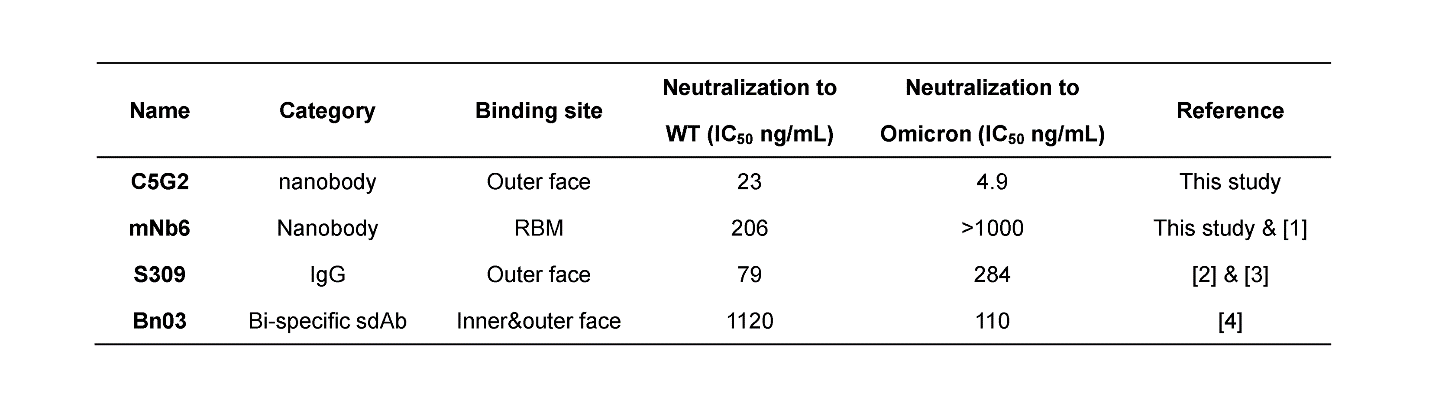

**Figure S5 Comparison of C5G2 binding to RBD with representative SARS­-CoV-2 neutralising antibodies.** The RBD was colored in grey white. The bindings of C5G2, mNb6(7KKL), S309(6WPS) and Bn03(7WHK) to RBD were shown as RBD kept in a fix orientation. VHH and VH of S309 were colored in red. VL of S309 was colored in cyan. The binding sites and neutralization efficacies of each antibody to both WT virus and Omicron were shown in the table below.
